# Pathological Oxidation of PTPN12 Underlies ABL1 Phosphorylation in HLRCC

**DOI:** 10.1101/296194

**Authors:** Yang Xu, Paul Taylor, Joshua Andrade, Beatrix Ueberheide, Brian Shuch, Peter M. Glazer, Ranjit S. Bindra, Michael F. Moran, Marston W. Linehan, Benjamin G. Neel

## Abstract

Hereditary Leiomyomatosis and Renal Cell Cancer (HLRCC) is an inherited cancer syndrome associated with a highly aggressive form of type 2 papillary renal cell carcinoma (PRCC). Germ line inactivating alterations in *Fumarate Hydratase* (*FH*) cause HLRCC, and result in elevated levels of reactive oxygen species (ROS). Recent work indicates that *FH -/-*PRCC cells have increased ABL1 activation, which promotes tumor growth, but how ABL1 is activated remained unclear. Oxidation can regulate protein-tyrosine phosphatase (PTP) catalytic activity; conceivably, ROS-catalyzed inactivation of an ABL-directed PTP might account for ABL1 activation in this malignancy. Previously, our group developed “q-oxPTPome,” a method that can globally monitor the oxidation of classical PTPs. We have now refined the q-oxPTPome approach, increasing its sensitivity by >10X. Applying q-oxPTPome to FH-deficient cell models shows that multiple PTPs are either highly oxidized (including PTPN12) or overexpressed. In general, highly oxidized PTPs were those that have relatively high sensitivity to exogenous H_2_O_2_. Most PTP oxidation in FH-deficient cells is reversible, although nearly 40% of PTPN13 is oxidized irreversibly to the sulfonic acid state. Using “substrate-trapping mutants”, we mapped PTPs to their putative substrates, and found that only PTPN12 could target ABL1. Furthermore, knockdown experiments identify PTPN12 as the major ABL1 phosphatase in HLRCC. Overall, our results show that ROS-induced PTPN12 oxidation accounts for ABL1 phosphorylation in HLRCC-associated PRCC, reveal a novel mechanism for inactivating a tumor suppressor gene product, and establish a direct link between pathological PTP oxidation and neoplastic disease.

## Introduction

Reactive oxygen species (ROS), including superoxide anion (O_2^-^_), hydrogen peroxide (H_2_o_2_), and hydroxyl radical (^•^OH), are not just toxic by-products from aerobic respiration, but can also act as second messengers in cellular signaling pathways (1). Protein-tyrosine phosphatases (PTPs) have emerged as major ROS targets under physiological and pathological conditions. The PTP superfamily is encoded by more than 100 genes, including 38 classical PTPs, which specifically dephosphorylate phosphotyrosine (2, 3). Classical PTPs can be further subdivided into non-transmembrane PTPs (NT-PTPs) and receptor-like PTPs (R-PTPs). Nearly all NT-PTPs have a single catalytic (PTP) domain, whereas most R-PTPs have two PTP domains (D1, D2), with D1 typically harboring the catalytic activity and D2 exerting a regulatory function. Classical PTPs are defined by a conserved signature motif (I/V)H<u>**C**</u>SAGXXR(S/T)G, containing an essential catalytic cysteine residue that is critical for dephosphorylation. The catalytic cysteine has a low pKa and exists as a thiolate anion (S^-^) at physiological pH, rendering it quite susceptible to oxidization. Upon exposure to ROS, the PTP thiolate anion can be oxidized to the sulfenic acid (S-OH) state, which is labile and immediately rearranges to a disulfide (S-S) or sulfenylamide (S-N) bond. These cysteine oxidation events reversibly inhibit PTP activity and protect the catalytic cysteine from irreversible oxidation to sulfinic acid (SO_2_H) or sulfonic acid (SO_3_H) states at higher ROS concentrations. ROS-induced PTP oxidation has been reported to occur in physiological and pathological cellular processes, including growth factor signaling, cytokine signaling, integrin signaling, diabetes, atherosclerosis, chronic inflammation and cancer (4).

Hereditary leiomyomatosis and renal cell carcinoma (HLRCC) is an inherited cancer syndrome in which affected individuals are at risk for the development of a highly aggressive, poor prognosis form of Type 2 papillary renal cell carcinoma (PRCC) (5–7). HLRCC is associated with germ line inactivating alterations in *Fumarate Hydratase* (*FH*), which encodes a TCA cycle enzyme (8). FH loss-of-function results in accumulation of fumarate, which leads to inactivation of the antioxidant glutathione (GSH) and persistent oxidative stress (9, 10). Consequently, HLRCC cells have elevated ROS levels despite of hyperactivation of the anti-oxidant protein NRF2 (11–13). A recent study demonstrated that FH deficiency-induced ROS indirectly stimulate ABL1 phosphorylation and activation, which, in turn, is critical for *FH -/-*PRCC growth and survival (14). However, the precise mechanism by which ROS causes increased ABL1 activation remained undefined.

We hypothesized that the elevated ROS in HLRCC-associated PRCC might induce oxidation and inactivation of a specific PTP that normally dephosphorylates ABL1. Several methods for detecting PTP oxidation have been developed, including the in-gel PTPase assay, alkylation agent-linked biotin assays, dimedone-based assays, and specific antibodies against oxidized forms of PTPs (15). Nevertheless, identifying endogenously oxidized cellular PTPs remains challenging. Previously, we developed a mass spectrometry (MS)-based method, q-oxPTPome, that can monitor oxidation of all catalytically active classical PTPs (16). We substantially refined the q-oxPTPome protocol, combined it with the filter-aided sample preparation (FASP) method (17), and increased its sensitivity by at least 10-fold. Using this approach, we found that several PTPs are highly oxidized in FH-deficient PRCC cell lines, and identified PTPN12 oxidation as the cause of increased ABL1 tyrosine phosphorylation in these cells, establishing a novel mechanism of tumorigenesis via oxidative inhibition of PTPN12.

## Results and Discussion

### Refined q-oxPTPome can sensitively detect H_2_O_2_-induced classical PTP oxidation

For q-oxPTPome, reduced PTPs (PTP-S-) are alkylated with N-ethylmaleimide (NEM; PTP-S-NEM) to render them unreactive to other modifications, while the reversibly oxidized PTPs (PTP-SOH or other rearrangements) are converted to their sulfonic acid forms (PTP-SO_3_H) via the sequential use of dithiothreitol (DTT) and pervanadate (PV). As a result, q-oxPTPome detects the combination of reversible oxidized PTPs and PTPs that exist basally in the sulfonic acid state. This approach can also be modified to detect either total PTP levels (qPTPome) or the basal sulfonic acid state only (PTP-SO_3_H), but does not monitor the levels of PTPs in the sulfinic acid (PTP-SO_2_H) state (Fig. 1A).

**Figure 1.**
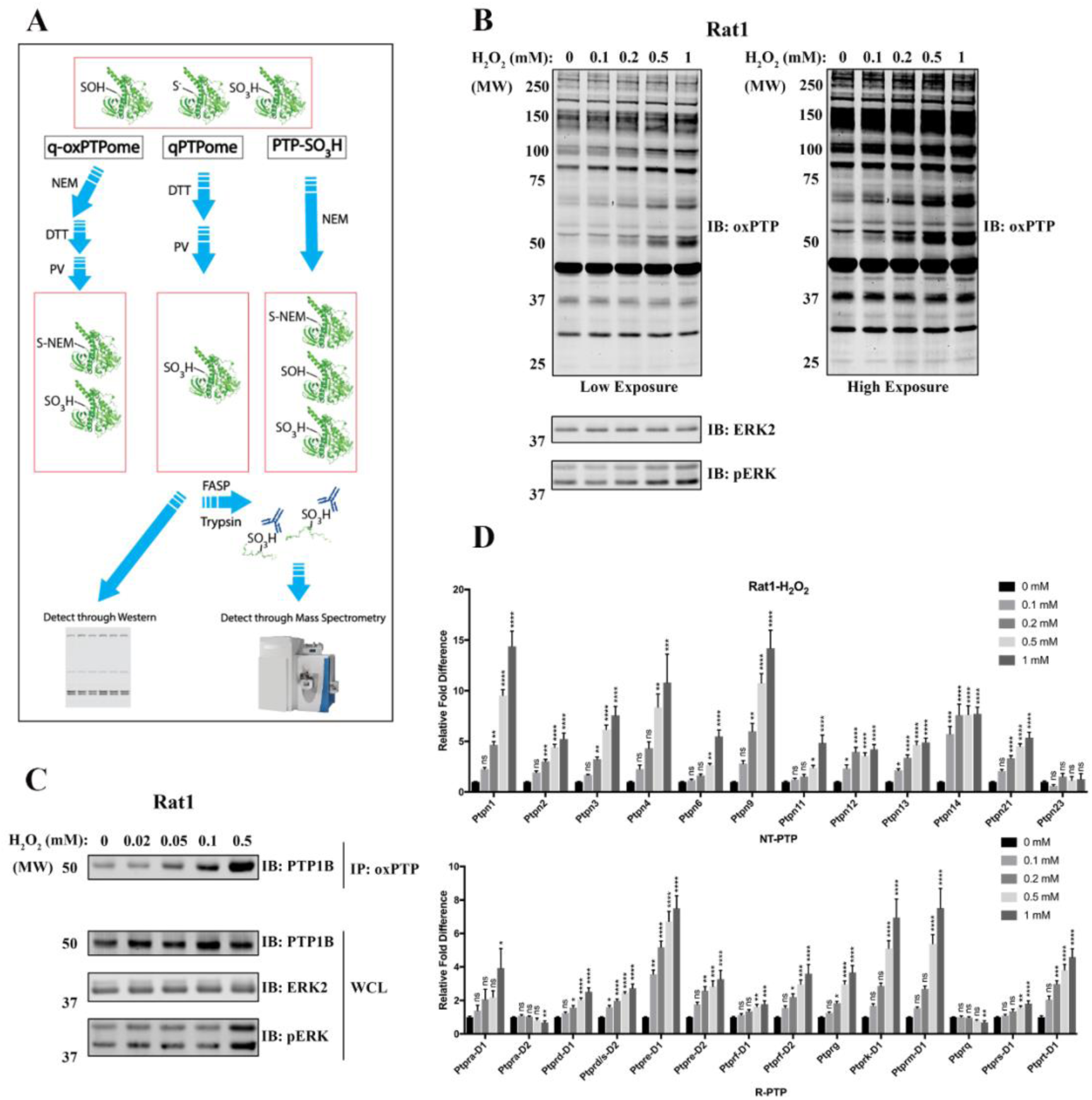
Refined q-oxPTPome can sensitively monitor H_2_o_2_-induced classical PTP oxidation. **A,** Scheme for q-oxPTPome, qPTPome and PTP-SO_3_H approaches; see text for details. NEM: N-ethylmaleimide, DTT: dithiothreitol, PV: pervanadate. FASP: filter-aided sample preparation. **B-D**, Rat1 cells were treated with H_2_O_2_ at the indicated concentrations for 4 min and then processed for q-oxPTPome. **B,** Samples were directly analyzed by immunoblotting using oxPTP Ab. **C,** Samples were immunoprecipitated using oxPTP Ab and immunoblotted for PTP1B (PTPN1). **D,** Samples were analyzed by mass spectrometry (MS) and quantified by MS label-free quantification. Data represent mean ± SEM (n=4-8; *p<0.05, **p<0.01, ***p<0.001, ****p<0.0001; ANOVA with Dunnett post-test).

The q-oxPTPome protocol was only capable of reproducibly quantifying PTP oxidation in cells exposed to extracellular H_2_o_2_ concentrations ≥1 mM (16). We further optimized the q-oxPTPome protocol and incorporated the FASP purification method (17) into our approach to increase the sensitivity (see Materials and Methods). To test our refined protocol, we treated Rat1 and HeLa cells with 0.1-1 mM of H_2_O_2_, subjected cell lysates to q-oxPTPome, and resolved the treated lysates by SDS-PAGE, followed by immunublotting with oxPTP monoclonal antibody (Ab). We observed multiple bands, whose intensities increased with increasing H_2_o_2_ levels, consistent with increased oxidation of these proteins (Fig. 1B and Supplementary Fig. S1A). To confirm that PTPs were oxidized, treated lysates were subjected to immunoprecipitation with the ox-PTP antibody, followed by immunoblotting for PTP1B (encoded by *Ptpn1*). As expected, PTP1B showed a dose-dependent increase in oxidation with increased H_2_o_2_ exposure, with levels as low as 50 nM capable of evoking detectable PTP1B oxidation (Fig. 1C, Supplementary Fig. S1C and Fig. S1D). Because q-oxPTPome detects both reversible oxidation and the irreversible, sulfonic acid state, we compared PTP-SO_3_H levels (Supplementary Fig. S1B, lane 1 and 2) with q-oxPTPome (Supplementary Fig. S1B, lane 3 and 4). As anticipated, nearly all of the detectable bands were in the q-oxPTPome lanes, consistent with reversible oxidation. We also transfected HEK293 cells with WT or catalytic cysteine-mutated (CS) Flag-*PTPN1* or Flag-*PTPN9* expression constructs, and assessed oxidation of the heterologously expressed PTPs. Only the cognate WT PTPs were detected under basal or H_2_O_2_-treated conditions, after immunoprecipitation with ant-Flag antibodies and immunoblotting with the oxPTP Ab (Supplementary Fig. S1E). These data confirm that the improved q-oxPTPome approach only detects PTP catalytic cysteine oxidation, and comport with our earlier report (16).

To assess oxidation of the entire classical PTP family, PTP oxidation induced by 0.1-1 mM H_2_o_2_ treatment of Rat1 or HeLa cells was quantified using MS (Fig. 1D and Supplementary Fig. S1F). We detected >20 classical PTPs, including the D1 and D2 domains of some R-PTPs, in each cell line. Most PTPs showed differential and gradually increasing levels of PTP oxidation in response to increasing concentrations of H_2_o_2_, and many had statistically significant increases in oxidation at 0.1 or 0.2 mM of H_2_o_2_. This level of sensitivity is at least 10-fold higher than seen by our original approach (16). Consistent with our previous results, some PTPs (e.g., PTPN23, PTPRA-D2) did not show increased oxidation or even appeared to have decreased oxidation at ^high concentrations of H_2_o_2_. These findings reflect oxidation of these PTPs to the sulfinic acid^ state, which cannot be detected by our assay (16). Our refined q-oxPTPome assay detects exogenous ROS-induced PTP oxidation at levels comparable to the most sensitive approaches in the literature (15). Furthermore, its ability to monitor oxidation of all classical PTPs simultaneously makes q-oxPTPome a useful tool to screen for oxidized PTPs in physiological and pathological states.

### Multiple PTPs are highly oxidized in FH-deficient cells

Because FH-deficient cell has enhanced ABL1 tyrosyl phosphorylation, which is indirectly the consequence of increased ROS levels (14), we suspected ROS-induced oxidation/inactivation of an ABL1-directed PTP. To begin to address thus possibility, we compared the UOK262 cell line, which is derived from an *FH*-mutant HLRCC patient (UOK262 FH-deficient) to UOK262 cells with wild-type *FH* re-introduced (UOK262 FH-WT) (18) or YUNK1 (Yale University Normal Kidney) cells, an SV40-immortalized kidney cell line with or without *FH* knockdown (YUNK1 and YUNK1 FH-KD). In accord with the previous studies, ROS levels were increased and ABL1 was hyper-phosphorylated in FH-deficient cells (Supplementary Fig. S2A and Fig. 2A). Using the refined q-oxPTPome or qPTPome approaches, we observed several proteins that exhibited increased oxidation in UOK262 FH-deficient cells, compared with their FH-reconstituted counterparts (q-oxPTPome; Fig. 2B left panels), without a change in their expression levels (qPTPome; Fig. 2B right panel). MS demonstrated a global increase in classical PTP oxidation, normalized to the expression level of each PTP, with PTPN7 (a.k.a. HePTP), PTPN9 (a.k.a. PTP-MEG2), PTPN12 (a.k.a. PTP-PEST) and PTPN13 (a.k.a. FAP-1 or PTP-BAS) being the top four oxidized PTPs (Fig. 2C). Increased PTPN12 oxidation was confirmed using immunoblotting (Fig. 2D and Supplementary Fig. S2D). Although most PTPs were expressed at similar levels in both cell lines, PTPN13 and PTPN18 (a.k.a. BDP1 or PTP-HSCF) appeared to be overexpressed in UOK262 FH-deficient, compared with FH-WT-reconstituted, cells (Supplementary Fig. S2B). The global increase in PTP oxidation is consistent with high oxidative stress in FH-deficient cells, suggesting the PTP catalytic cysteines being sensitive ROS targets. In addition, UOK262 FH-WT cells were treated with H_2_o_2_ and different PTPs had various oxidative sensitivity (Supplementary Fig. S2C). Notably, PTPN7, PTPN9 and PTPN12, the top three PTPs oxidized in response to H_2_o_2_, were also the most highly oxidized in FH-deficient cells (Fig. 2C and Supplementary Fig. S2C), implying that PTPs that are more sensitive to ROS might be more vulnerable to FH deficiency-induced oxidative stress.

**Figure 2.**
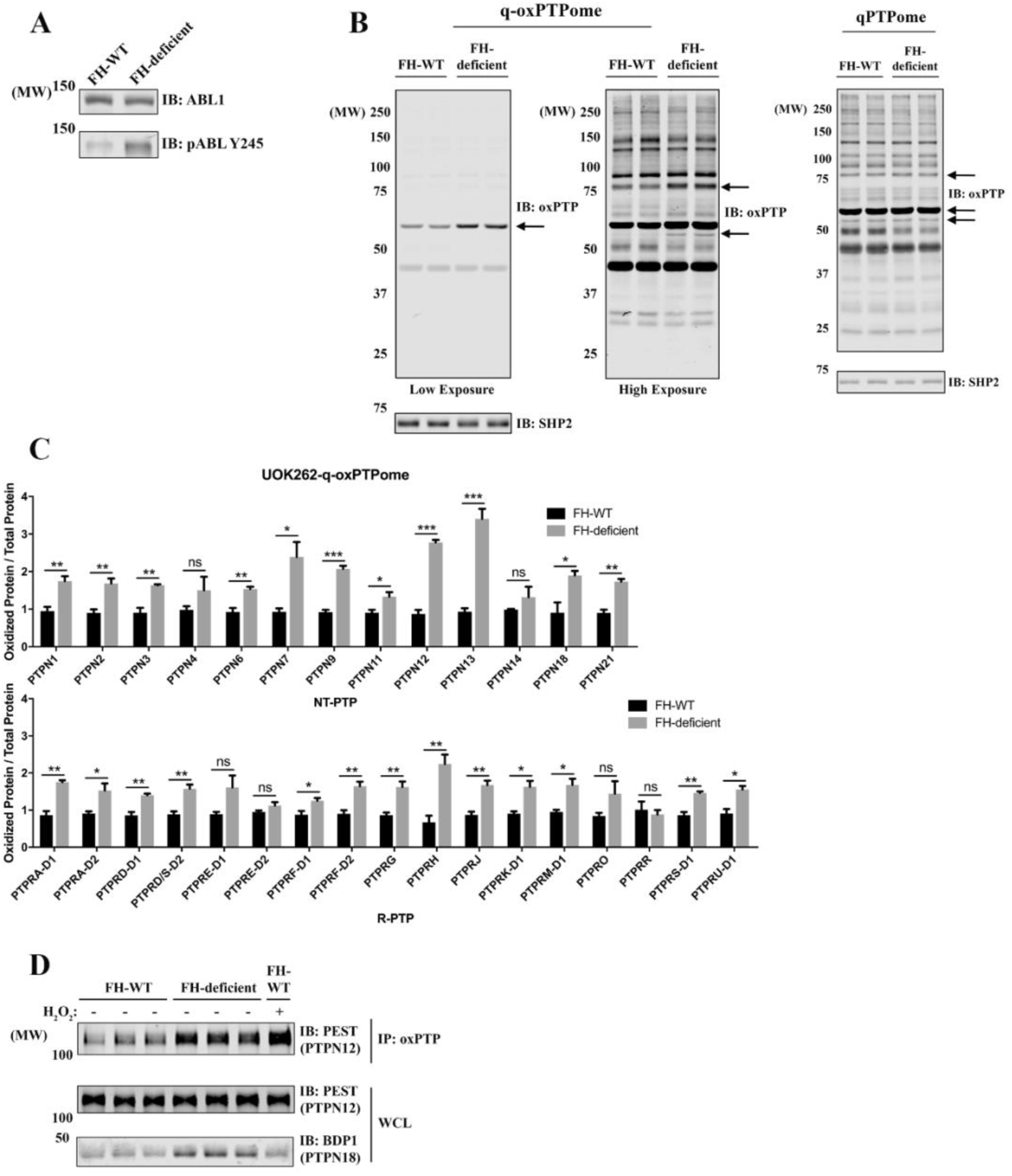
Multiple classical PTPs are oxidized in FH-deficient PRCC cell lines. **A,** UOK262 (FH-WT and FH-deficient) cells were lysed, and the levels of ABL1 and pABL-Y245 were assessed by immunoblotting. **B,** UOK262 (FH-WT or FH-deficient) cells were processed by q-oxPTPome (left panels) or qPTPome (right panel), and immunoblotted with oxPTP Ab. **C,** Samples from Fig. 2B were analyzed by MS and label-free quantification. The q-oxPTPome signal was normalized to the qPTPome PTP signal (to adjust for PTP expression level). Data represent mean ± SEM (n=3; *p<0.05, **p<0.01, ***p<0.001, ****p<0.0001; unpaired two-tailed t test). **D,** UOK262 (FH-WT or FH-deficient) were processed through q-oxPTPome. Samples were immunoprecipitated with oxPTP Ab and immunoblotted for PTPN12/PEST. H_2_o_2_, 1 mM for 4 min.

Because q-oxPTPome detects both reversible oxidation and the irreversible, sulfonic acid state of PTPs, we asked how much of the observed oxidation was due to the latter. As shown in Supplementary Fig. S2E, several proteins had increased SO_3_H hyper-oxidation in FH-deficient cells, which was also seen when we treated cells with high concentrations (10 mM) of H_2_o_2_. Although we only loaded 1/3 of the q-oxPTPome samples compared with other samples (Supplementary Fig. S2E last lane vs. the rest), most bands from the q-oxPTPome lane (alkylated, reduced, terminally oxidized and then immunoblotted with the oxPTP antibody) had a much higher signal intensity than in the SO_3_H hyper-oxidation lanes (alkylated and then immunoblotted with the oxPTP antibody), indicating that most PTP (or protein-thiol) oxidation in FH-deficient cells was reversible. This result was supported by MS analysis, shown as the percentage of SO_3_H hyper-oxidation, relative to q-oxPTPome signal from FH-deficient cells (Supplementary Fig. S2F). Most PTPs had low or undetectable SO_3_H hyper-oxidation (Supplementary Fig. S2F), except for PTPN4 and PTPN13, which exhibited 30% and 40% oxidation to the SO_3_H state, respectively. This finding implies that these PTPs are either intrinsically more vulnerable to irreversible oxidation or they localize in a cellular compartment(s) that has(ve) higher levels of FH deficiency-induced ROS.

### PTPN12 trapping mutant can interact with phosphorylated ABL1

Because PTPN7, PTPN9 and PTPN12 have the highest oxidation in FH-deficient cells without overexpression (Fig. 2C and Supplementary Fig. S2C), we asked whether any of these PTPs can regulate ABL1 phosphorylation. The PEST-type PTP subfamily, including PTPN12, PTPN18 and PTPN22, has been reported to dephosphorylate ABL1 (19). Because PTPN18 is expressed in all tested HLRCC cell lines and is even overexpressed in UOK262 FH-deficient cells (Fig. 2D and Supplementary Fig. S2B), we included PTPN18 in this analysis.

PTP substrate-trapping mutants, including mutants of the catalytic cysteine to serine (CS) or the aspartate in the WPD motif to alanine (DA), have been used widely for substrate identification; such mutants often retain substrate binding capacity without catalyzing dephosphorylation (20). We generated HEK293 cells that inducibly express WT flag-tagged *PTPN7*, *PTPN9*, *PTPN12* and *PTPN18* or their cognate CSDA mutants, in response to tetracycline. These cells were pre-treated with or without pervanadate, which can further boost the intracellular phosphotyrosine by inactivating all cysteine-based PTPs. As shown in Supplementary Fig. S3A and S3B, after pervanadate pre-treatment, only cells expressing PTPN12-CSDA interacted with a 140 kDa protein (consistent with the expected size of ABL1) on phosphotyrosine blots. Using both in-gel and gel-free MS affinity purification, we found that only the PTPN12 co-immunoprecipitated (co-IPed) ABL1, together with other known PTPN12 substrates, such as p130Cas and PSTPIP2 (Fig. 3A, Supplementary Fig. S3D, S3E, and S3F). Immunoblotting confirmed that PTPN12-CSDA interacts with ABL1 under non-denaturing detergent conditions but not in an SDS-containing buffer (Supplementary Fig. S3C), consistent with a typical trapping mutant/substrate interaction. Similar results were obtained using HeLa cells (Fig. 3B).

**Figure 3.**
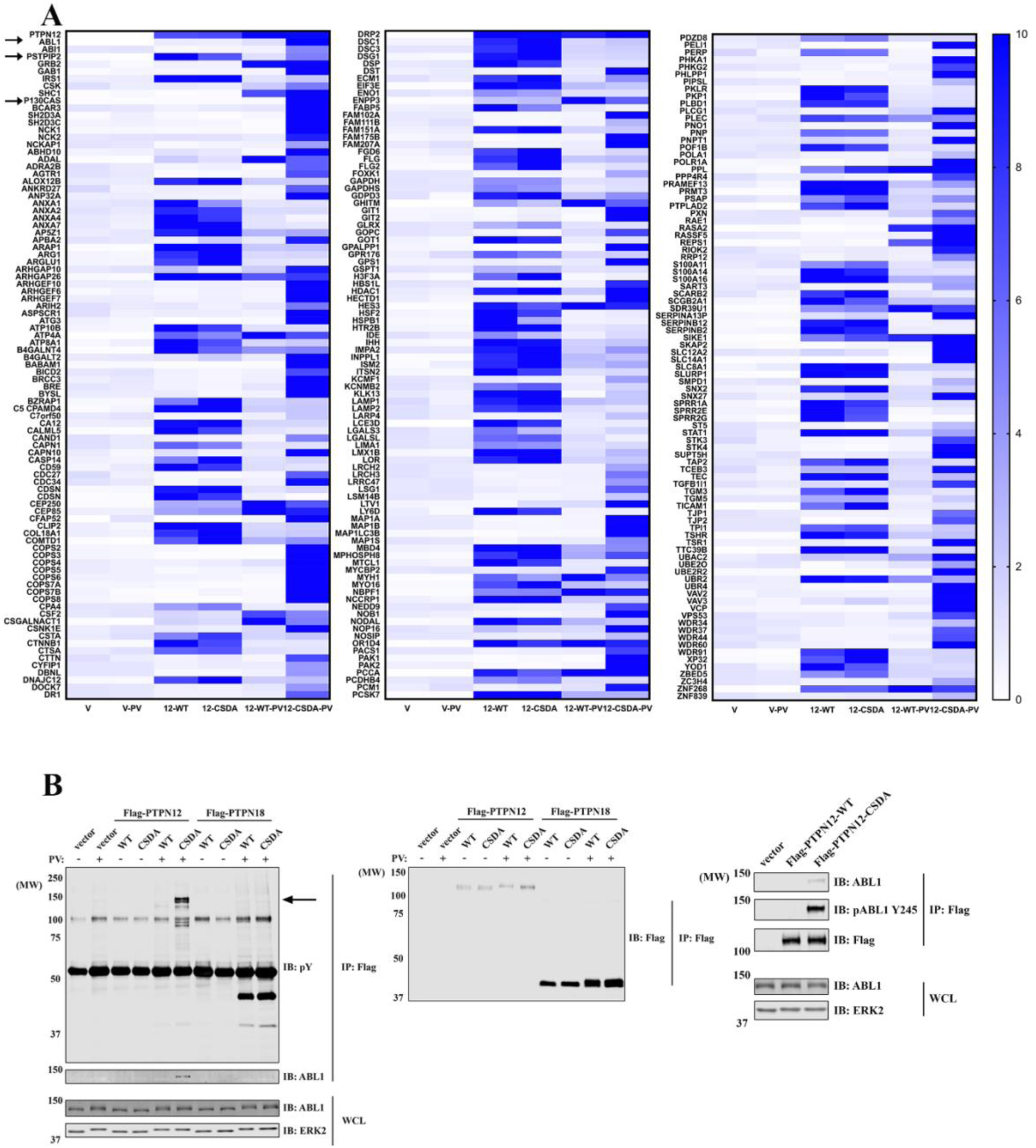
PTPN12/PEST, but not PTPN18/BDP1, can trap tyrosyl phosphorylated ABL1. **A,** HEK293 T-REx cells, treated with tetracycline to induce expression of Flag vector (V), Flag-*PTPN12*-WT, *Flag-PTPN12*-CSDA, were exposed to 50 μM PV or buffer control for 30 min before lysis. PTP-interacting proteins were co-immunoprecipitated with Flag beads and identified by MS (combining results of both gel-free affinity purification and in-gel trypsin digestion). Heat maps show the relative amounts of each protein identified (compared with Flag vector control samples) as determined by label-free quantification. **B,** HeLa T-REx cells, treated with tetracycline to induced expression of Flag vector, Flag-*PTPN12*-WT, Flag-*PTPN12*-CSDA, Flag-*PTPN18*-WT, or Flag-*PTPN18*-CSDA, were exposed to 50 μM PV or buffer control for 30 min before lysis. Samples were immunoprecipitated with Flag beads, and immunoblotted with ABL1, pABL-Y245, and Flag Abs.

PEST-type PTPs were reported to dephosphorylate ABL1 via the adaptor PSTPIP1 (19). Surprisingly, however, we only observed the related adaptor PSTPIP2 in PTPN12 or PTPN18 immunoprecipitates (Fig. 3A and Supplementary Fig. S3F). PSTPIP2 lacks the critical SH3 domain that interacts with ABL1 to bridge it to PEST-type PTPs (Supplementary Fig. S4A), we hypothesized that failure of PTPN18 to interact with ABL1 might be due to the low or no expression of PSTPIP1 in the tested cell lines. To test this hypothesis, we overexpressed PSTPIP1 in HeLa cells, and consistent with previous results, PSTPIP1 co-IPed with PTPN12 and PTPN18 (Supplementary Fig. S4B). Furthermore, PSTPIP1 expression helped PTPN18 to interact with ABL1. However, unlike ABL1 recovered in PTPN12-CSDA immunoprecipiates, ABL1 that co-IPed with PTPN18 was not heavily tyrosine-phosphorylated (Supplementary Fig. S4B), suggesting that PTPN12 might be a better ABL1 phosphatase regardless whether or not PSTPIP1 is expressed.

### PTPN12 regulates ABL1 phosphorylation in *FH -/-*PRCC cell lines

To investigate the function of PEST-type PTPs in *FH -/-*PRCC cell lines, we knocked down *PTPN12* or *PTPN18* in UOK262 FH-WT or FH-deficient cell lines. *PTPN12*-KD, but not *PTPN18*-KD, cells showed increased phosphorylation of a 140 kDa protein in total phosphotyrosine immunoblots, as well as increased tyrosine-phosphorylated ABL1, as detected by a pY-specific antibody (Fig. 4A left panel). Similar results were obtained using another HLRCC patient-derived *FH -/-*PRCC cell line, UOK268 (Fig. 4A right panel). We then knocked down ABL1 and found the intensity of PTPN12 KD-induced hyper-phosphorylated 140 kDa band was markedly decreased (Fig. 4B), indicating that most, if not all, of hyper-phosphorylated 140 kDa species to be ABL1.

**Figure 4.**
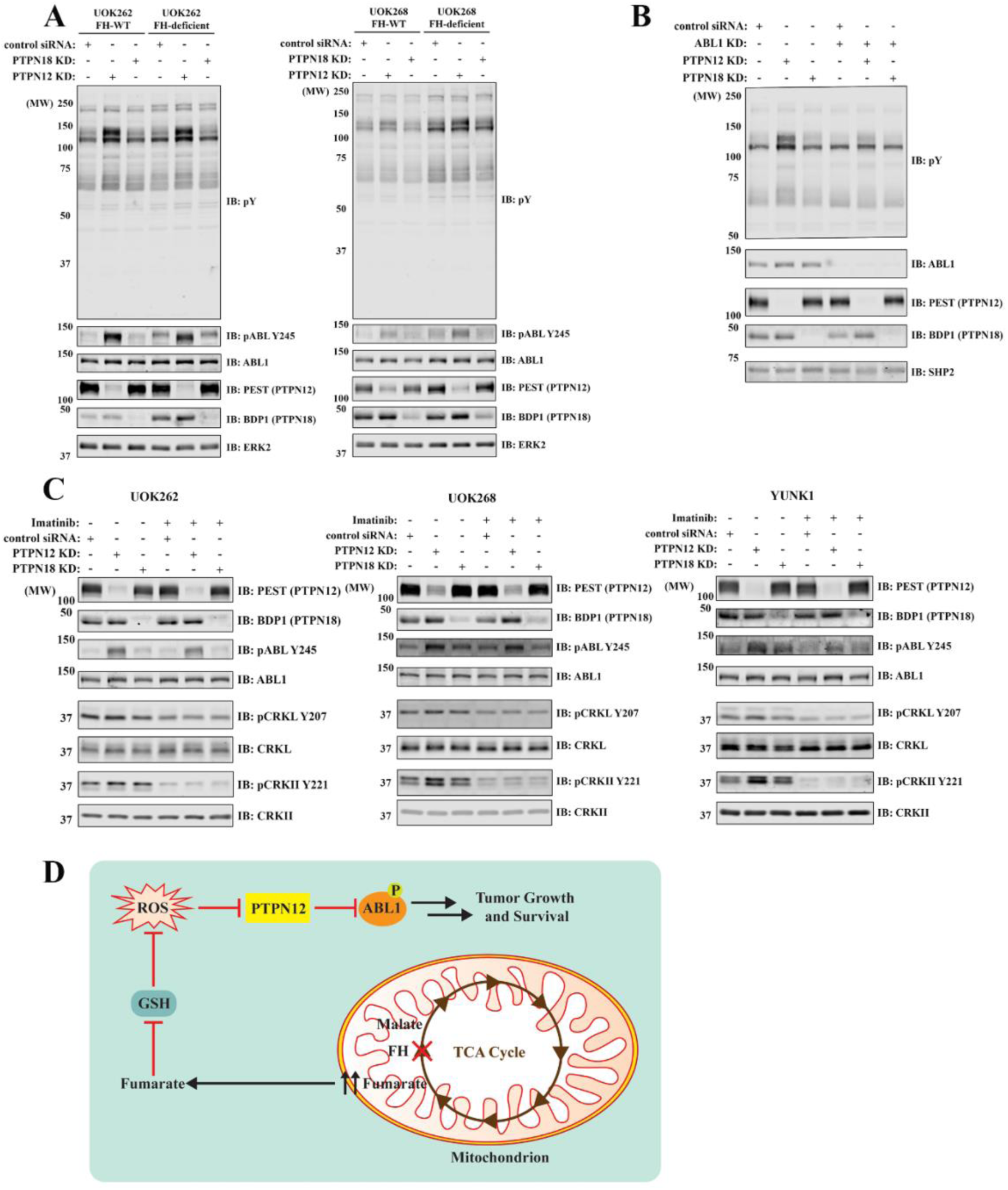
PTPN12/PEST regulates ABL1 phosphorylation and downstream signaling in FH-deficient PRCC cell lines. **A,** UOK262 (FH-WT and FH-deficient) and UOK268 (FH-WT and FH-deficient) cells were treated with control, *PTPN12* or *PTPN18* siRNAs before lysis and immunoblotting with pY or pABL-Y245 Abs. **B,** UOK262 FH-WT cells were treated with control, *ABL1*, *PTPN12* or *PTPN18* siRNAs before lysis and immunoblotting with anti-pY Ab. **C,** UOK262 FH-WT, UOK268 FH-WT or YUNK1 cells were treated with control, *PTPN12* or *PTPN18* siRNAs and then with or without 2 μM imatinib for 1 hr before lysis and immunoblotting with the indicated antibodies. **D,** Schematic showing how FH deficiency-induced ROS oxidize/inhibit PTPN12, which leads to ABL1 hyper-phosphorylation and FH-deficient PRCC tumor growth and survival. GSH: glutathione.

We investigated also assessed ABL1-dependent downstream signaling pathways. CRKII was hyper-phosphorylated in UOK268 and YUNK1 cells after *PTPN12* KD, but there was not much difference in UOK262 cells (Fig. 4C). Increased CRKII phosphorylation was reversed by the Imatinib, consistent with ABL1 being the kinase acting on CRKII. However, after PTPN18-KD, there is a slightly increase in ABL1 and CRKII phosphorylation in YUNK1 cells (Fig. 4C right panel): this difference cannot be explained by PSTPIP1 because all of the cell lines have undetectable levels of PSTPIP1 by IB and qPCR (Supplementary Fig. S4C and S4D). Therefore, whereas PTPN12 is the major PTP regulating ABL1 phosphorylation in FH-deficient cells, other PEST-type PTPs might play a minor role. Because multiple other classical PTPs were oxidized in FH-deficient cells (Fig. 2C), we cannot rule out the possibility that they might also control ABL1 phosphorylation.

In conclusion, our refined q-oxPTPome approach can sensitively and globally quantify classical PTP oxidation, and allowed us to discover that PTPN12 is highly oxidized in FH-deficient HLRCC-associated PRCC cell lines as a consequence of their elevated oxidative stress. We found PTPN12 can trap ABL1 in the absence of adaptor protein PSTPIP1 and is the major PTPs regulating ABL1 phosphorylation in HLRCC cell lines. Because ABL1 phosphorylation/activation is critical for FH-deficient PRCC cells to proliferate and survive (14), *PTPN12* appears to be an important, although “dynamically regulated” tumor suppressor gene in HLRCC (Fig. 4D). Previous studies showed that PTPN12 can suppress mammary epithelial cell proliferation and transformation and is an important tumor suppressor in triple-negative breast cancer (TNBC). In TNBC, however, PTPN12 is mainly inactivated through deletion or loss-of-function mutations (21). Our data demonstrate that oxidative stress-induced PTPN12 oxidation can inhibit PTPN12 without deletion/mutation, and suggest that activation of (a) PTK(s) via ROS-mediated inactivation of cognate PTPs is a novel mechanism for cancer pathogenesis.

## Materials and Methods

### Cell Lines and Cell Culture

HEK293 T-REx cells were obtained from ThermoFisher Scientific. HeLa T-REx cells were kindly provided by Dr. Brian Raught (Princess Margaret Cancer Centre, Canada). UOK262 FH-WT, UOK262 FH-deficient, UOK268 FH-WT and UOK268 FH-deficient cell lines were kindly provided by Dr. Marston Linehan (National Cancer Institute, Bethesda, USA). YUNK1 and YUNK1 FH-KD cells were kindly provided by Dr. Ranjit Bindra (Yale Therapeutic Radiology, USA). HepG2 and THP-1 cells were purchased from ATCC. THP-1 cells were cultured in RPMI-1640, while all other cells were cultured in Dulbecco’s modified Eagle’s medium (DMEM). All media contained 10% FBS (Corning) and 100 Units/ml penicillin/streptomycin (Gibco).

### Immunoblotting

Cells were lysed in a buffer containing 50 mM HEPES (pH 7.5), 150 mM NaCl, 2 mM EDTA, 1% NP40, 0.1% SDS with a protease and phosphatase inhibitor cocktail (40 mg/ml PMSF, 2 mg/ml antipain, 2 mg/ml pepstatin A, 20 mg/ml leupeptin, and 20 mg/ml aprotinin, 20 mM NaF, 10mM β-glycerophosphate and 2 mM Na_3_VO_4_). Homogenates were centrifuged at 16,100 × g for 15 min at 4 °C and supernatants were collected. Total protein of 10-50 µg was resolved by SDS-PAGE and analyzed by immunoblotting. Antibodies used for immunoblots included: oxidized PTP active site antibody (oxPTP, clone 335636) and anti-mouse PTP1B antibody from R&D Systems; anti-ERK2 antibody (D-2), anti-SH-PTP2 antibodies (C-18 and B1) and anti-ABL1 antibody (8E9) from Santa Cruz Biotechnology; anti-Flag antibodies from Sigma; anti-PTPN12 antibody from Bethyl; anti-PSTPIP1 antibody from Abcam; anti-phosphotyrosine antibody (4G10) from EMD Millipore; anti-pERK antibody (9101), anti-c-ABL antibody (2862), anti-phospho-ABL1 (Tyr245) antibody, anti-PTPN18 antibody, anti-phospho-CRKL (Tyr207) antibody, anti-CRKL antibody, anti-phospho-CrkII (Tyr221) antibody and anti-CrkII antibody from Cell Signaling Technology. siRNAs for PTPN12, PTPN18, ABL1 were purchased from Dharmacon.

### q-oxPTPome

A lysis buffer containing 20 mM Tris-HCl (pH 7.35), 100 mM NaCl, 10% glycerol, 1% NP40) was degassed overnight with vacuum at RT, and then transferred to a hypoxia workstation (Don Whitley Scientific). Immediately before using, 30 mM N-ethylmaleimide (Sigma), 0.5 U/µL catalase (Calbiochem), 0.125 U/mL superoxide dismutase (Sigma) and a protease inhibitor cocktail (2 mg/ml antipain, 2 mg/ml pepstatin A, 20 mg/ml leupeptin, and 20 mg/ml aprotinin) were added to the lysis buffer. Within the hypoxia workstation, cells were quickly washed once with degassed PBS, immediately lysed in the above buffer, and incubated for 1 hr at 4°C in the dark with constant rotation. Homogenates were clarified at 10,000 × g for 1 min at 4°C. Supernatants were collected in the hypoxia workstation and buffer exchanged by gel filtration (GE Healthcare Life Sciences) into degassed solution (20 mM HEPES [pH 7.4], 100 mM NaCl, 10% glycerol, 0.05% NP40, 0.02 U/µL catalase). Desalted lysates were treated with 5 mM DTT and rotated in the dark for 1.5 hrs at 37°C. Within the hypoxia workstation, the lysates were buffer exchanged again to remove DTT and freshly prepared pervanadate was added to a final concentration of 8.5 mM. The samples were then incubated overnight at 4°C in the dark with constant rotation. The next day, 4 mM EDTA was added to the solution, and the samples were ready to proceed to immunoblotting, immunoprecipitation or MS (See Supplemental material for detailed methods).

### Quantitative Real-Time PCR

RNA was extracted using the miRNeasy Mini Kit (Qiagen), reverse transcribed and analyzed by Sybr green-based qPCR (Sigma), according to the manufacturer’s instructions. The forward and reverse primers for PSTPIP1 were: GCA GCA TAG ACG CCG ACA TC and CTT CCG TGC AGC AGT CCA G, respectively (22), while the forward and reverse primers for GAPDH were: GAG TCA ACG GAT TTG GTC GT and TTG ATT TTG GAG GGA TCT CG, respectively.

### ROS Measurement

Cells were treated as indicated, incubated 30 min with 500 nM DCF-DA (Invitrogen) at 37°C in phenol red-free media, trypsinized, washed with cold PBS, and analyzed by flow cytometry (CytoFlex).

### Statistical Analysis

Data are presented as the mean ± SEM. Statistical significance was determined using unpaired Student’s t test or one-way ANOVA (comparing all the experimental groups to the same basal control group), as appropriate. If ANOVA was significant, individual differences were evaluated using the Dunnett post-test. Statistical analyses were performed using Graph-Pad Prism 7, with p < 0.05 considered significant.

## Acknowledgements

The authors thank Dr. Anne Marie Pendergast (Duke University Medical School) for pointing out the hyperactivation and requirement for ABL1 in HLRCC. We thank Drs. Anne-Claude Gingras (Lunenfeld-Tanenbaum Research Institute) and Nicole St-Denis (University of Toronto) for providing various WT PTP constructs. We thank Drs. Yubao Wang and Kwan-Ho Tang (NYU Langone) for help with flow cytometry and Dr. Jieyu Jen (NYU Langone) for assistance with qPCR. This work was supported by R01CA49152 (BGN), the Proteomics Shared Resource of the Perlmutter Cancer Center (P30 CA16087), and the Princess Margaret Cancer Foundation. MFM is supported by Canada Research Chairs, Leukemia and Lymphoma Society of Canada, Canadian Institutes of Health Research, and Canadian Cancer Society. The mass spectrometric experiments were in part supported by NYU School of Medicine, the Laura and Isaac Perlmutter Cancer Center Support grant P30CA016087 from the National Cancer Institute and a shared instrumentation grant from the NIH, 1S10OD010582-01A1 for the purchase of an Orbitrap Fusion Lumos.

